# ORIGAMI: Orientation-Aware Graph Neural Network for Assessing Multimeric Interfaces of Protein Complex Structures

**DOI:** 10.64898/2026.05.31.729128

**Authors:** Xinyu Wang, Debswapna Bhattacharya

**Affiliations:** Department of Computer Science, Virginia Tech, Blacksburg, Virginia, 24061, USA

## Abstract

Deep learning-based protein structure prediction methods have led to a paradigm-shift in computational structural biology, yet reliably assessing the quality of computationally predicted multimeric structures remains challenging. Recent methods have demonstrated benefits of employing graph neural networks for assessing multimeric interfaces of protein complexes, but ignore geometric orientational features naturally occurring in 3-dimensional protein conformational space and act only on scalar weights. We present ORIGAMI, an orientation-aware graph neural network for assessing multimeric interfaces of protein complex structures that leverages both scalar and 3D vector node representations to perform symmetry-aware geometric operations while maintaining SO(3)-equivariance by capturing fine-grained orientational relationships between residues across protein-protein interfaces to estimate the interface Local Distance Difference Test (iLDDT) score. Tested on targets from multiple rounds of Critical Assessment of Structure Prediction (CASP) challenges, ORIGAMI achieves superior performance across multiple interface quality assessment benchmarks, with particularly strong gains in the expanded CASP16 interface-level evaluation and in controlled comparisons against both non-equivariant and equivariant graph neural network baselines. It also demonstrates robust cross-metric generalization by reproducing superposition-based DockQ scores with high fidelity, despite being trained only to estimate the superposition-free iLDDT score. ORIGAMI is freely available at https://github.com/Bhattacharya-Lab/ORIGAMI.

## 1 Introduction

Proteins perform essential biological functions, from structural support to enzyme catalysis, making them central to therapeutic applications. Recent breakthroughs in structure prediction, exemplified by AlphaFold3 (1), RFDiffusion (2), and Boltz-2 (3), have revolutionized structural modeling capabilities (4). Understanding protein-protein interactions in multimeric complexes is essential for elucidating biological mechanisms (5), identifying therapeutic targets (6), and advancing drug design (7). However, while AI-driven tools excel at modeling individual protein domains, accurately predicting binding affinities and interaction strengths in multimeric complexes remains challenging (8; 9). The reliability of computationally predicted multimeric structures varies considerably due to system complexity, sequence quality, and algorithmic constraints (10; 11). Consequently, developing Estimation of Model Accuracy (EMA) methods specifically for multimeric proteins has become imperative for establishing confidence in predicted interactions and guiding experimental validation (12; 13). The dramatic improvement in interface contact prediction from 31% in CASP14 to 90% in CASP15 (14) demonstrates that interface-centric evaluation strategies offer a promising approach for selecting optimal multimeric models.

While experimental techniques like X-ray crystallography and cryo-electron microscopy can reveal protein complex structures, their high resource demands have driven the development of computational modeling methods (15). Current methodologies for estimating protein complex quality can be broadly categorized into three approaches: biophysics-based, statistics-based, and machine-learning-based methods. Biophysics-based methods rely on fundamental physical principles to evaluate binding interactions. These include molecular dynamics simulations for binding free energy calculations, such as evERdock (16), force field-based approaches including AMBER (17) and CHARMM (18), and continuum solvation methods like MM/PBSA and MM/GBSA (19; 20). Statistics-based methods derive scoring functions from statistical analysis of known protein structures. Examples include knowledge-based functions such as DFIRE (21), ITScore (22), GOAP (23), PMF (24), DrugScore (25), and RAPDF (26), which extract energy estimates from normalized occurrence frequencies of atomic contacts in structural databases. Machine-learning-based methods represent the newest category, employing artificial intelligence to learn complex patterns from structural and sequence data. These methods incorporate sophisticated architectures such as graph neural networks and transformers, as demonstrated by VoroIF-GNN (27), DeepRank-GNN-esm (28), DProQA (29), GATE (30), GNN-DOVE (31), EquiRank (32) and PIsToN (33). Machine learning approaches can effectively integrate outputs from both biophysics-based and statistics-based methods as input features, enabling more comprehensive structural representations. This hybrid approach has demonstrated superior performance in recent Critical Assessment of protein Structure Prediction competitions (34; 35), highlighting the advantage of combining multiple methodological paradigms for enhanced protein complex quality assessment.

Despite these advances, two critical limitations persist in current approaches. First, many existing interface quality assessment methods rely primarily on scalar residue, contact, or voxel-level representations, which limits their ability to explicitly model orientation-dependent interface geometry. Although geometric deep learning architectures such as Geometric Vector Perceptron(GVP) (36), E(3)-equivariant graph neural networks (EGNN) (37), Tensor Field Networks (TFN) (38), SE(3)-Transformer (39), and Equiformer (40) have introduced vector- or tensor-valued representations for molecular modeling, their use has not been systematically explored for multimeric interface quality assessment. The inherent complexity of protein-protein interactions and the diverse structural features of multimeric assemblies demand specialized neural network architectures capable of comprehensively modeling protein complex geometry—encompassing both backbone and side-chain conformations across multiple interacting chains. Recent advances demonstrate that orientation-based architectures with directed weight designs significantly enhance protein quality assessment accuracy (41). However, these approaches remain limited to single-chain protein quality assessment, raising a critical question: can vector features and directed weight designs be leveraged to assess protein multimeric interfaces? To address this gap, ORIGAMI adapts orientation-aware scalar–vector message passing, directed vector-valued weights, and cross-product geometric filters for interface quality estimation of multimeric protein complexes. Second, most current quality assessment methods target superposition-based metrics as ground truth labels (28; 31; 30; 29; 32; 33), requiring structural alignment with native structures, a procedure that is not always optimal. Using superposition-based metrics for multimeric interfaces introduces a fundamental anomaly: scores depend on overall structural alignment rather than interface-specific quality, potentially yielding inaccurate estimates (42). This limitation prompts another critical question: can superposition-free metrics better assess protein multimeric interfaces?

Interface LDDT (iLDDT) offers a promising solution to this superposition dependence. As a superposition-free metric, iLDDT (1; 43) evaluates protein-protein interface accuracy by measuring local distance differences between atoms across interface chains, assessing how well inter-chain atomic interactions are reproduced in predicted structures. As an all-atom variant of lDDT (43) that focuses exclusively on interface contacts, iLDDT provides atomic-level precision ideal for interface quality assessment. These considerations underscore the need for specifically designed neural network architectures that effectively capture essential geometric features while maintaining robustness to disordered regions and non-participating accessory domains. Motivated by these challenges, we introduce ORIGAMI, a novel orientation-aware deep graph learning method that targets iLDDT scores as ground truth labels for protein complex evaluation for the first time. ORIGAMI extends orientation-aware graph neural networks by transforming individual weights from scalars to 3D vectors, enabling enhanced representation of critical geometric features for multimeric protein interfaces (41).

ORIGAMI extracts the interface region from multimeric structures, constructs a graph focused exclusively on the residues within the extracted interface, and outputs a single global accuracy score for the interface. We evaluated ORIGAMI’s performance using targets from multiple rounds of Critical Assessment of Structure Prediction (CASP) challenges, including CASP15 (14) and CASP16 (35). Experimental results demonstrate that ORIGAMI consistently outperforms state-of-the-art methods, including the top-performing CASP predictors, across a wide range of evaluation metrics. Notably, ORIGAMI exhibits remarkable robustness by reproducing superposition-based DockQ scores with high fidelity, despite being trained exclusively on the superposition-free iLDDT metric. An open-source software implementation of ORIGAMI, licensed under the GNU General Public License v3, is freely available at https://github.com/Bhattacharya-Lab/ORIGAMI.

## 2 Materials and Methods

### 2.1 Overview of the ORIGAMI Architecture

Protein–protein interfaces are governed not only by residue identity and proximity, but also by directional geometric cues such as side-chain orientation, local surface curvature, and backbone frame alignment. These anisotropic patterns are critical for binding specificity and stability, yet are difficult to capture with conventional scalar-only graph neural networks (GNNs), which primarily encode distances and scalar descriptors. ORIGAMI addresses this limitation by introducing an orientation-aware graph neural architecture that represents interfacial residues using paired scalar–vector features and processes them with directed weight operators. In contrast to standard scalar-parameterized layers, these operators treat network parameters as 3D vectors, enabling the model to express directional filters that respond to specific geometric configurations (e.g., aligned, orthogonal, or twisted orientations). All geometric computations are performed within residue-specific local frames, ensuring strict SE(3)-equivariance throughout the network. The full architecture consists of six stacked Directed Weight Perceptron (DWP) layers (Fig. 1c) integrated into an equivariant message-passing framework (Fig. 1a). The depth of six layers was selected based on an ablation study on the VoroIFGNN test dataset (Supplementary Section 2.1), in which we varied the number of DWP layers and observed that six provided the best trade-off between predictive performance and computational cost. Each layer refines scalar biochemical descriptors and vector geometric representations. After the final layer, node features are aggregated via scatter mean pooling and processed by a lightweight readout network composed of linear layers, ReLU activations, and dropout, yielding a scalar prediction corresponding to the interface Local Distance Difference Test (iLDDT) score in the range [0, 1] (43; 1). Architectural specifications are provided in Supplementary Material 1.4.

**Figure 1.**
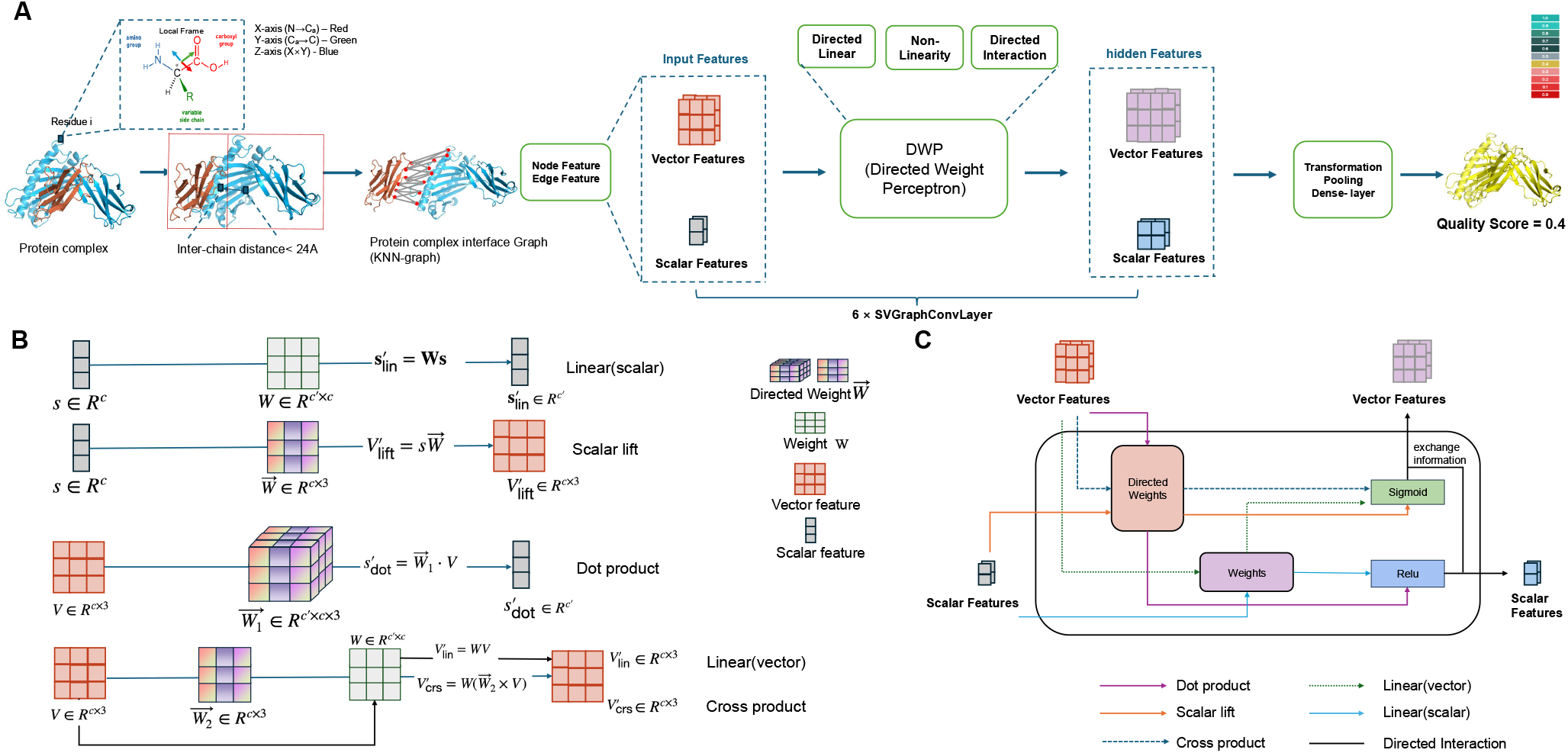
**A,** Overview of the ORIGAMI framework. **B**, Directed Linear Module, the first component of the Directed Weight Perceptron, transforms input features through geometric operations: 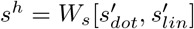 and 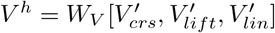, enabling rich scalar-vector feature interactions. **C**, Directed Weight Perceptron.

### 2.2 Graph Construction and Feature Representation

#### Spatial k-nearest neighbor interface graph

For each predicted complex, ORIGAMI constructs a spatial *k*-nearest neighbor (k-NN) graph over interfacial residues. We identify interfacial residues as those containing any atom within a 24 Å cutoff from an atom in a different chain. This radius reflects the optimal balance observed in our ablation experiments (Supplementary Section 2.1), where we evaluated distance thresholds and found 24 Å to retain biologically relevant inter-chain contacts while minimizing noise from distal residues. Inter-chain interactions at interfaces are dominated by short-range geometric and physicochemical effects; thus, a local neighborhood graph provides a faithful approximation of the interaction surface. Each interface is represented as a graph *G* = (*V, E*), where *V* denotes interfacial residues and *E* contains edges connecting each residue to its *k* nearest neighbors in ℝ^3^. Consistent with our ablation study on the VoroIFGNN test dataset (Supplementary Section 2.1), we set *k* = 40, which preserved over 97% of true interfacial contacts across CASP complexes while avoiding excessive graph density. Formally, for each node *u* ∈*V*, we connect *u* to its 40 nearest neighbors by Euclidean distance, generating a spatial k-NN graph that emphasizes local geometric context while filtering distant, potentially noisy interactions (44; 45; 46; 47).

#### Scalar–vector node and edge featurization

ORIGAMI employs a dual-modality representation for both nodes and edges (see **Table 1**). Each residue node *u* ∈*V* is characterized by scalar features **s**_*u*_ ∈ℝ^33^ and vector features **V**_*u*_ ∈ℝ^3*×*3^. Scalar components encode amino acid identity, secondary structure, solvent accessibility, backbone torsion angles, and additional physicochemical descriptors reflecting the local biochemical environment. Vector components represent residue-level orientation via local backbone and side-chain directions, allowing the model to capture alignment-dependent cues that are invisible to scalar-only representations. Each edge (*i, j*) ∈*E* is associated with scalar features **s**_*ij*_ ∈ℝ^34^ and vector features **V**_*ij*_ ∈ℝ^1*×*3^. Scalar edge features include sequence separation, inter-residue distances and distance transforms, contact-type indicators, and geometric descriptors of the local interface geometry. Vector edge features consist of normalized displacement vectors between residues, encoding the direction of approach across the interface. Together, these scalar–vector features enable ORIGAMI to jointly model biochemical properties and orientation-dependent geometric relationships that govern interface complementarity and binding stability.

**Table 1.**
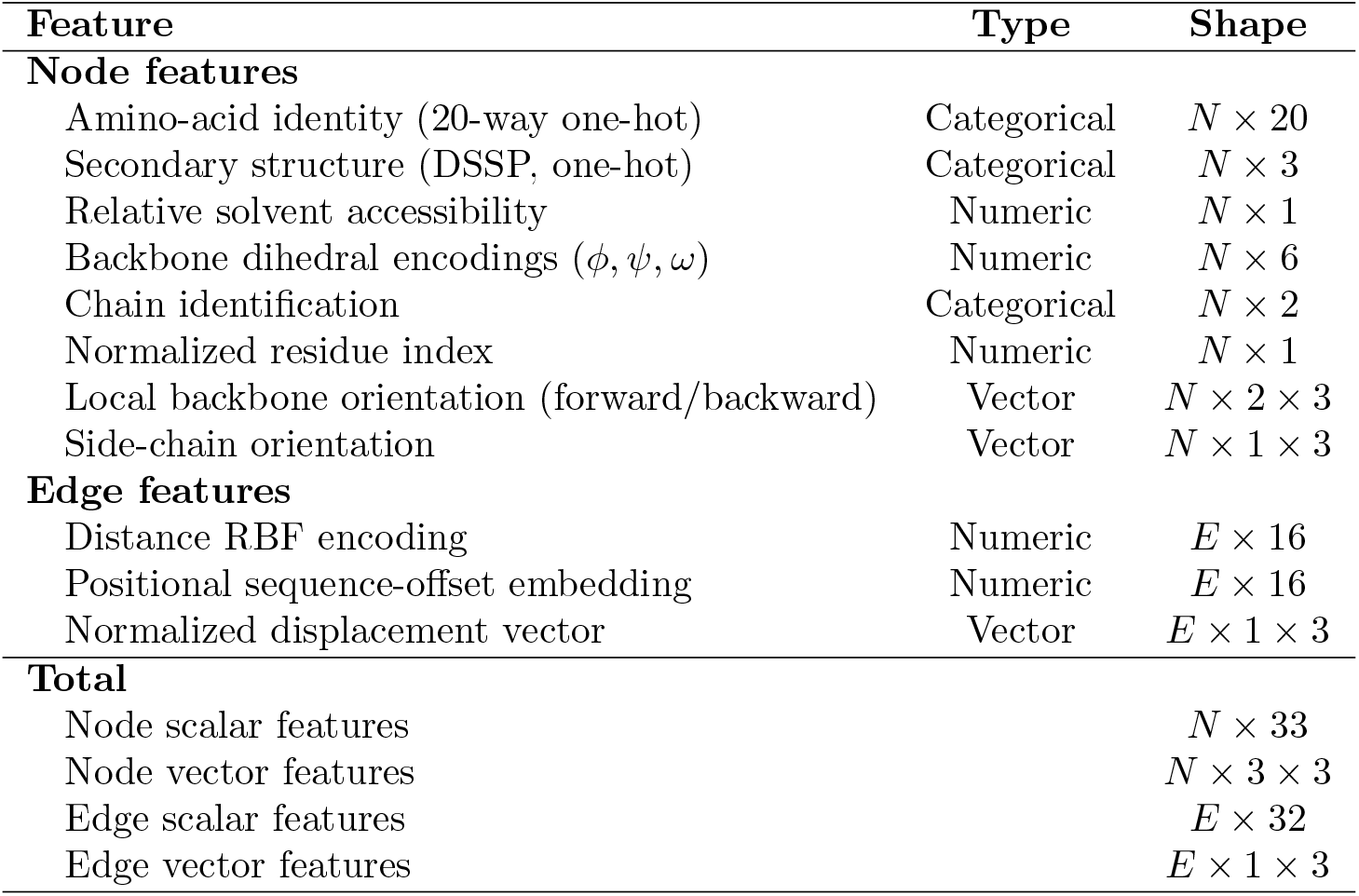
Summary of ORIGAMI node and edge features. **Bold** indicates the best performance; underlined indicates the second-best performance for each metric.

| Feature | Type | Shape |
| --- | --- | --- |
| **Node features** |  |  |
| Amino-acid identity (20-way one-hot) | Categorical | $N \times 20$ |
| Secondary structure (DSSP, one-hot) | Categorical | $N \times 3$ |
| Relative solvent accessibility | Numeric | $N \times 1$ |
| Backbone dihedral encodings $(\phi, \psi, \omega)$ | Numeric | $N \times 6$ |
| Chain identification | Categorical | $N \times 2$ |
| Normalized residue index | Numeric | $N \times 1$ |
| Local backbone orientation (forward/backward) | Vector | $N \times 2 \times 3$ |
| Side-chain orientation | Vector | $N \times 1 \times 3$ |
| **Edge features** |  |  |
| Distance RBF encoding | Numeric | $E \times 16$ |
| Positional sequence-offset embedding | Numeric | $E \times 16$ |
| Normalized displacement vector | Vector | $E \times 1 \times 3$ |
| **Total** |  |  |
| Node scalar features | | $N \times 33$ |
| Node vector features | | $N \times 3 \times 3$ |
| Edge scalar features | | $E \times 32$ |
| Edge vector features | | $E \times 1 \times 3$ |

### 2.3 Directed Weight Perceptron

Conventional GNNs for protein complex assessment typically parameterize network weights as scalars and apply them to scalar inputs. Even when vector features are used, scalar weights generally act only through norm scaling or linear projections, limiting the model’s ability to learn direction-selective filters. To overcome this limitation, ORIGAMI introduces the Directed Weight Perceptron (DWP) (Fig. 1c), a learnable SE(3)-equivariant module parameterized by vector-valued weights

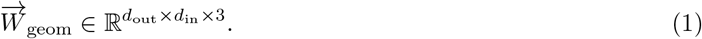

Each directed weight encodes a learnable geometric direction that can be aligned, contrasted, or combined with residue-level vector features.

A DWP layer operates on scalar–vector tuples (*s, V*) and consists of three components: a directed linear module (Fig. 1b), non-linear modulation, and scalar–vector interaction.

#### Directed linear module

The module applies four geometric operators to (*s, V*):

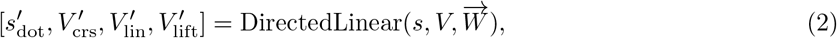

where *s* and *V* denote input scalar and vector features.

#### Dot-product filters

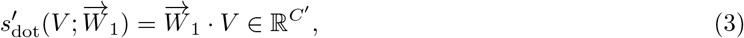

which measures alignment between residue orientations and learned directional weights, enabling detection of specific orientational patterns.

#### Cross-product filters

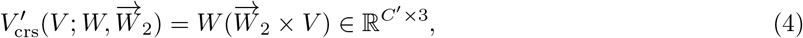

which generates vectors perpendicular to the plane spanned by *V* and 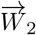, capturing orthogonal and rotational geometric relationships.

#### Linear vector transforms

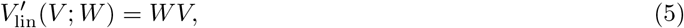

corresponding to standard vector-space mappings.

#### Scalar-to-vector lifting

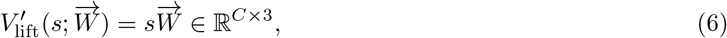

which maps scalar biochemical signals into directional space, allowing the network to associate residue-level scalar properties with geometric orientations.

These four operators together form an expressive, strictly SE(3)-equivariant geometric basis that subsumes scalar-weighted and norm-only vector parameterizations, enabling anisotropic geometric reasoning within interfacial environments.

#### Non-linearity module

The intermediate outputs 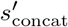 and 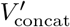, obtained by concatenating the directed linear outputs, are passed to a non-linear transformation:

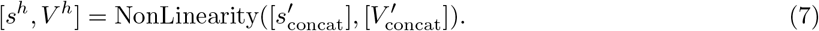

Scalar components *s*^*h*^ are processed with ReLU activations to introduce non-linearity while maintaining numerical stability. Vector components *V* ^*h*^ are modulated by a magnitude-aware gating function that preserves direction:

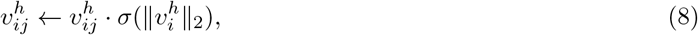

where *σ* denotes the sigmoid function. This gating adjusts vector magnitudes based on confidence while retaining orientation, which is crucial for maintaining SE(3)-equivariance and preserving biologically meaningful geometric information.

#### Scalar–vector interaction module

Finally, the scalar and vector streams are coupled through a directed interaction module:

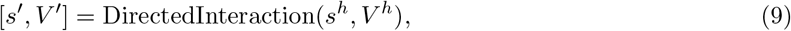

which integrates scalar biochemical descriptors with vector geometric features via learned weighted combinations. This bidirectional coupling allows scalar features to modulate geometric filters (e.g., conditioning on residue type or local environment) and allows geometric patterns to influence scalar representations, yielding a unified representation of interface quality that jointly reflects chemical and spatial information.

### 2.4 Equivariant Message Passing Protocol

To ensure SE(3)-equivariance across all layers, ORIGAMI performs message passing and DWP updates within residue-specific local coordinate systems. For each residue *u*, we construct an orthonormal local frame using backbone atoms 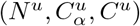 (Fig. 1a):

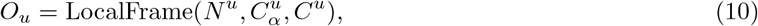

where *O*_*u*_ ∈ℝ^3*×*3^ is an orthogonal matrix whose columns define the local axes. All vector features associated with residue *u* and its incident edges are transformed into this frame prior to processing.

Message passing at layer *l* proceeds via

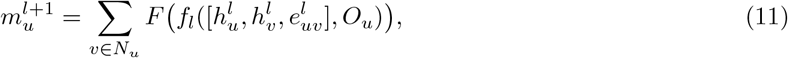

where *N*_*u*_ denotes the neighborhood of *u*, 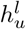 and 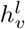 are scalar–vector node features at layer *l*, 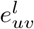are edge features, and *F* and *f*_*l*_ are learned transformations that operate in the local frame defined by *O*_*u*_. Node features are then updated as

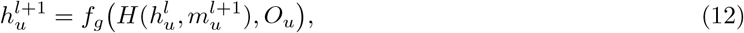

where *H* combines previous node features and aggregated messages, and *f*_*g*_ applies the final DWP-based transformation in the local coordinate system before mapping back to global coordinates. Because all vector operations (dot product, cross product, and linear transforms) are SE(3)-equivariant, and all coordinate changes are orthogonal, the overall architecture preserves strict equivariance under global rigid-body transformations while remaining computationally efficient with message-passing complexity 𝒪 (|*V* |*k*).

### 2.5 Estimation of Interface Local Distance Difference Test

The accuracy of protein–protein interfaces is evaluated using the interface Local Distance Difference Test (iLDDT), a superposition-free metric quantifying how well local distance relationships are preserved between atoms across different chains (43; 1). For each atom pair (*i, j*) belonging to different chains and within an inclusion radius *R*_0_ = 15 Å, we define the distance difference

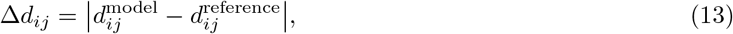

where 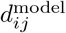and 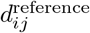are the inter-atomic distances in the predicted model and reference structure, respectively. Distance preservation is assessed at four tolerance thresholds *t*_1_ = 0.5 Å, *t*_2_ = 1.0 Å, *t*_3_ = 2.0 Å, and *t*_4_ = 4.0 Å.

The iLDDT score is computed as

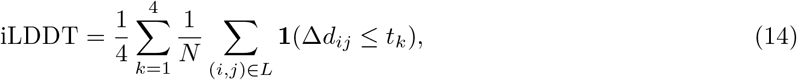

where *L* is the set of inter-chain atom pairs within *R*_0_, *N* = |*L*|is the total number of such pairs, and **1**(·) is the indicator function. The resulting iLDDT score lies in [0, 1], with higher values indicating better preservation of interfacial geometry. A detailed description of the implementation is provided in Supplementary Material 1.3.

### 2.6 Training Objective and Implementation Details

ORIGAMI is trained with a multi-objective loss combining a single-structure regression term and a pairwise ranking term. The regression term encourages accurate prediction of individual interface quality scores, while the pairwise term preserves the relative ordering of decoys within each target, which is essential for selecting high-quality protein complex models.

For each protein complex model *i*, ORIGAMI predicts an iLDDT score 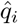, with the corresponding ground-truth score denoted by *q*_*i*_. The single-structure regression loss is defined using the Huber loss:

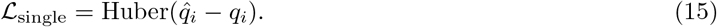

To promote correct ranking of decoys from the same target, we additionally use a pairwise ranking loss. For a pair of models (*i, j*) belonging to the same target, the predicted and true score differences are defined as

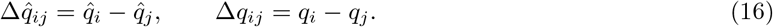

The pairwise loss is then given by

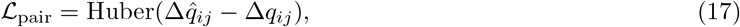

which penalizes discrepancies in relative model quality and discourages inversions in the predicted ranking within each target.

The complete training objective combines the two components:

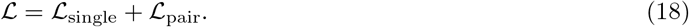

This dual-objective formulation enables ORIGAMI to predict accurate individual iLDDT scores while maintaining proper relative rankings among decoys, supporting both protein complex quality assessment and comparative model selection. We optimized the network with Adam using a learning rate of 10^*−*5^ and dropout of 0.15 for 200 epochs on 8 NVIDIA A100 GPUs with 80 GB memory. Despite using vector-weighted SE(3)-equivariant layers, the architecture remains computationally efficient and trains on large datasets with standard GPU resources.

### 2.7 Datasets, Benchmarking, and Evaluation Metrics

#### Voro-CASP datasets

We constructed benchmark datasets by combining protein complex models generated by VoroIFGNN (27) with structures from the CASP13 (48) and CASP14 (49) experiments. For each target, candidate models were selected using a two-stage protocol based on VoroMQA-energy (50) and interface CAD-score (51), yielding up to 20 models per target (12.8 on average) spanning a broad range of interface and binding-site accuracies. The resulting dataset contains 1,541 targets and 20,703 models, partitioned into training (1,090 targets, 15,279 models), validation (224 targets, 2,671 models), and testing (227 targets, 2,753 models) with no target overlap. Detailed dataset construction procedures are provided in Supplementary Material 1.2.

#### CASP15 and CASP16 datasets

We further evaluated ORIGAMI on recent CASP benchmarks by collecting predictions from the CASP16 and CASP15 websites (35). For CASP16, we evaluated 39 multimeric targets, including both dimeric and higher-order oligomeric assemblies, using reference-defined contacting subunit-pair interfaces. Interacting subunit pairs were identified from the experimental assemblies, and all submitted models were evaluated on the same set of contacting interfaces for each target, yielding 60,314 model–interface instances from 12,916 models. We additionally considered a CASP16 dimeric subset consisting of the common set of dimeric protein complexes predicted by the five top-performing predictors, comprising 12 targets and 3,000 models. For CASP15, we evaluated 24 dimeric targets and 5,264 models from 10 top-performing predictors and the three baseline methods.

#### Evaluation metrics

We evaluated model performance using Pearson correlation (*r*) for linear accuracy, Spearman correlation (*ρ*) for ranking consistency, and mean absolute error (MAE) for absolute prediction accuracy. We additionally reported the hit rate—the proportion of targets with at least one high-quality model (iLDDT ≥ 0.236) in the top-*N* predictions—and ROC-AUC using thresholds of iLDDT ≥ 0.236 and DockQ ≥ 0.23 (1). Complete metric definitions are provided in Supplementary Material 1.6.

## 3 Results

### 3.1 Performance evaluation on the CASP16 datasets

#### Interface-level evaluation on 39 CASP16 multimeric targets

We evaluated ORIGAMI on 39 CASP16 multimeric targets, including both dimeric and higher-order oligomeric assemblies, using reference-defined contacting subunit-pair interfaces. For each target, interacting subunit pairs were identified from the experimental assembly, and all submitted models were evaluated on the same set of contacting interfaces, yielding 60,314 model–interface instances from 12,916 models. **Table 2** presents the global performance comparison between ORIGAMI and the baseline methods, where all model–interface instances were pooled into a single evaluation set. ORIGAMI achieves the best performance across all reported metrics under both iLDDT and DockQ evaluation. For iLDDT, ORIGAMI obtains Pearson/Spearman/MAE values of 0.527/0.471/0.155, substantially outperforming VoroIF-GNN, DProQA, and DeepRank-GNN. For DockQ, ORIGAMI similarly achieves the strongest performance, with Pearson/Spearman/MAE values of 0.455/0.449/0.158. These results indicate that ORIGAMI can accurately evaluate multimeric interface quality not only in dimers, but also in interfaces embedded within higher-order CASP16 assemblies.

**Table 2.** Performance comparison of ORIGAMI and baseline methods on 39 CASP16 targets, including both dimeric and higher-order multimeric assemblies, using reference-defined contacting subunit-pair interfaces.

| Methods | iLDDT |  |  | DockQ |  |  |
| --- | --- | --- | --- | --- | --- | --- |
| | $r \uparrow$ | $\rho \uparrow$ | MAE ↓ | $r \uparrow$ | $\rho \uparrow$ | MAE ↓ |
| **ORIGAMI** | **0.527** | **0.471** | **0.155** | **0.455** | **0.449** | **0.158** |
| VoroIF-GNN | <u>0.382</u> | <u>0.359</u> | 0.360 | <u>0.330</u> | <u>0.322</u> | 0.396 |
| DProQA | 0.192 | -0.081 | <u>0.189</u> | 0.231 | -0.040 | <u>0.167</u> |
| DeepRank-GNN-ESM | 0.293 | 0.309 | 0.249 | 0.234 | 0.264 | 0.247 |

**Bold** indicates the best performance; underlined indicates the second-best performance for each metric.

Although the global evaluation provides a useful summary of overall predictive performance, it may obscure target-to-target or interface-to-interface variability and can be influenced by targets or interfaces with more submitted models. To further examine whether the global performance remains consistent across individual targets and interfaces, we conducted per-target and per-interface analyses. For the per-target analysis, all model–interface instances from the same CASP16 target were pooled, and one Spearman correlation was computed for each target. For the per-interface analysis, one Spearman correlation was computed separately for each reference-defined contacting subunit-pair interface across submitted models. **Figure 2** presents the resulting distributions, which reflect not only the average ranking performance of each method, but also its variability and stability across different targets and interfaces.

**Figure 2.**
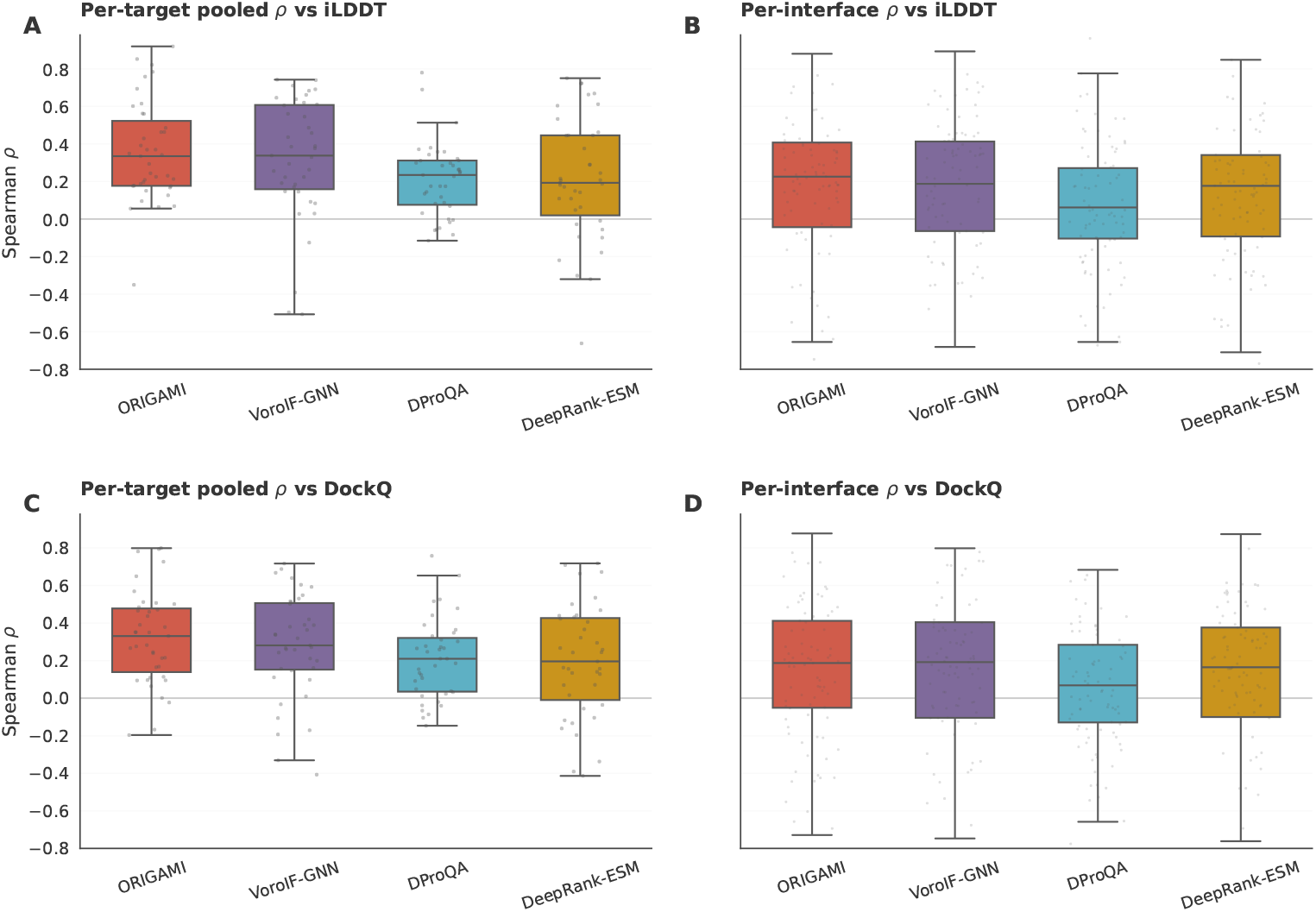
Benchmark comparison with distributions across 39 CASP16 targets and interfaces. **(A, C)** Per-target pooled Spearman correlations between predicted interface scores and reference scores for iLDDT and DockQ, respectively. **(B, D)** Per-interface Spearman correlations for iLDDT and DockQ, respectively. Boxes show the interquartile range, center lines show medians, whiskers show 1.5 × IQR, and points indicate individual target- or interface-level correlations.

For per-target pooled correlations against iLDDT, ORIGAMI achieves a mean Spearman correlation of 0.356 ± 0.266, better than VoroIF-GNN (0.319 ±0.325) and much higher than DProQA (0.210 ±0.198) and DeepRank-GNN-ESM (0.212 ±0.325). A similar trend is observed against DockQ, where ORIGAMI achieves the highest mean Spearman correlation of 0.322 ± 0.248, followed by VoroIF-GNN (0.279 ±0.277), DProQA (0.206 ±0.212), and DeepRank-GNN-ESM (0.197± 0.298). The per-interface analyses show the same overall trend, although ORIGAMI and VoroIF-GNN are closely matched, particularly for DockQ. Overall, these results complement the global evaluation by demonstrating that ORIGAMI maintains competitive performance when target-level and interface-level variability are considered.

#### Generalization to AlphaFold-generated CASP16 models

To directly evaluate whether ORIGAMI generalizes beyond the Voro-CASP training distribution, we assessed its model-selection performance on CASP16 AlphaFold2 and AlphaFold3 models. Specifically, the AlphaFold2 models were obtained from the CASP16 predictor group (#145), while the AlphaFold3 models were obtained from the CASP16 predictor group (#304). We report the mean per-target ranking loss, defined as the difference between the best available DockQ/iLDDT score among all decoys and the DockQ/iLDDT score of the top-ranked decoy selected by each method. Therefore, a lower ranking loss indicates that the method selects a model closer to the oracle-best decoy for each target. As shown in Table 3, ORIGAMI achieves the lowest ranking loss across both AlphaFold2 and AlphaFold3 model sets under both iLDDT and DockQ evaluation. In particular, ORIGAMI obtains ranking losses of 0.023 and 0.021 for iLDDT on AlphaFold2 and AlphaFold3 models, respectively, and 0.024 and 0.025 for DockQ. The second-best method varies across settings, whereas ORIGAMI consistently maintains top performance in all four comparisons. These results suggest that, despite being trained on an older CASP-derived dataset with potential pre-selection bias, ORIGAMI retains strong model-selection ability on newer AlphaFold-generated decoys with different quality distributions.

**Table 3.** Mean ranking loss of ORIGAMI and competing methods on AlphaFold2 and AlphaFold3 models.

| Methods | iLDDT |  | DockQ |  |
| --- | --- | --- | --- | --- |
|  | AF2 Models ↓ | AF3 Models ↓ | AF2 Models ↓ | AF3 Models ↓ |
| **ORIGAMI** | **0.023** | **0.021** | **0.024** | **0.025** |
| VoroIF-GNN | <u>0.024</u> | 0.054 | <u>0.033</u> | 0.056 |
| DProQA | 0.037 | 0.033 | 0.047 | <u>0.033</u> |
| DeepRank-GNN-ESM | 0.037 | <u>0.032</u> | 0.050 | 0.034 |

Values report mean ranking loss. Lower is better. **Bold** indicates the best performance; underlined indicates the second-best performance for each column.

#### Comparison with top CASP16 single-model EMA groups

We collected predictions from five top-performing CASP16 single-model predictors (35) in the EMA category from the CASP16 website. Unlike the 39-target interface-level evaluation above, CASP16 EMA groups provide a single quality score for each submitted model rather than separate scores for individual subunit-pair interfaces. Therefore, we considered the common set of dimeric targets in this experiment for which model-level scores from all five predictors could be directly compared with ORIGAMI. As such, the evaluations were conducted on the common set of 12 targets with available experimental structures. iLDDT and DockQ scores were computed using OpenStructure (52) on the intersection of dimeric protein complex models predicted by all five predictors, as our analysis focuses exclusively on dimeric complexes and therefore does not include the full set of CASP16 targets.

**Table 4** presents the performance comparison between ORIGAMI and the top-performing predictors from CASP16. ORIGAMI achieves the highest Pearson correlation of 0.778 with respect to iLDDT, surpassing VifChartreuseJaune (0.722), VifChartreuse (0.603), and AF unmasked (0.656). With respect to DockQ, ORIGAMI maintains the highest Pearson correlation of 0.729, demonstrating consistent performance across both metrics. ORIGAMI also achieves the lowest MAE values of 0.174 with respect to iLDDT and 0.191 with respect to DockQ, indicating superior prediction accuracy. While AF unmasked shows slightly higher Spearman correlations of 0.805 and 0.802 with respect to iLDDT and DockQ, respectively, ORIGAMI’s lower MAE indicates more accurate absolute predictions.

**Table 4.** Performance comparison of ORIGAMI and top CASP16 single-model EMA groups on 12 dimeric targets with available experimental structures.

| Methods | iLDDT |  |  | DockQ |  |  |
| --- | --- | --- | --- | --- | --- | --- |
| | $r \uparrow$ | $\rho \uparrow$ | MAE ↓ | $r \uparrow$ | $\rho \uparrow$ | MAE ↓ |
| **ORIGAMI** | **0.778** | <u>0.760</u> | **0.174** | **0.729** | <u>0.693</u> | **0.191** |
| VifChartreuseJaune | <u>0.722</u> | 0.712 | <u>0.217</u> | <u>0.684</u> | 0.608 | 0.252 |
| PIEFold_human | -0.151 | 0.028 | 0.318 | -0.143 | -0.055 | 0.349 |
| GuijunLab-Pathreader | 0.256 | 0.590 | 0.341 | 0.297 | 0.532 | 0.380 |
| VifChartreuse | 0.603 | 0.686 | 0.241 | 0.663 | 0.672 | <u>0.222</u> |
| AF_unmasked | 0.656 | **0.805** | 0.259 | 0.649 | **0.802** | 0.307 |

**Bold** indicates the best performance; underlined indicates the second-best performance for each metric.

To comprehensively evaluate the discriminative power of top-performing CASP16 predictors and ORIGAMI, we conducted ROC curve analysis using the AlphaFold3 acceptability criteria (1), where acceptable models are defined as those with iLDDT *>* 0.236 or DockQ *>* 0.23. **Figure 3** presents the ROC curves comparing ORIGAMI with the top 5 single-model CASP predictors from CASP16 on the same 12 targets. ORIGAMI achieves the highest AUC of 0.930 with respect to iLDDT, surpassing AF unmasked (0.911), VifChartreuseJaune (0.889), and VifChartreuse (0.850), while substantially outperforming GuijunLab-PAthreader (0.699) and PIEFold human (0.405). With respect to DockQ, ORIGAMI achieves the highest AUC of 0.925, outper-forming AF unmasked (0.916), VifChartreuseJaune (0.880), VifChartreuse (0.851), GuijunLab-PAthreader (0.715), and PIEFold human (0.413). The ROC curves reveal that ORIGAMI consistently maintains high true positive rates across different false positive rate thresholds, particularly in the low false positive rate region where discriminative power is most critical.

**Figure 3.**
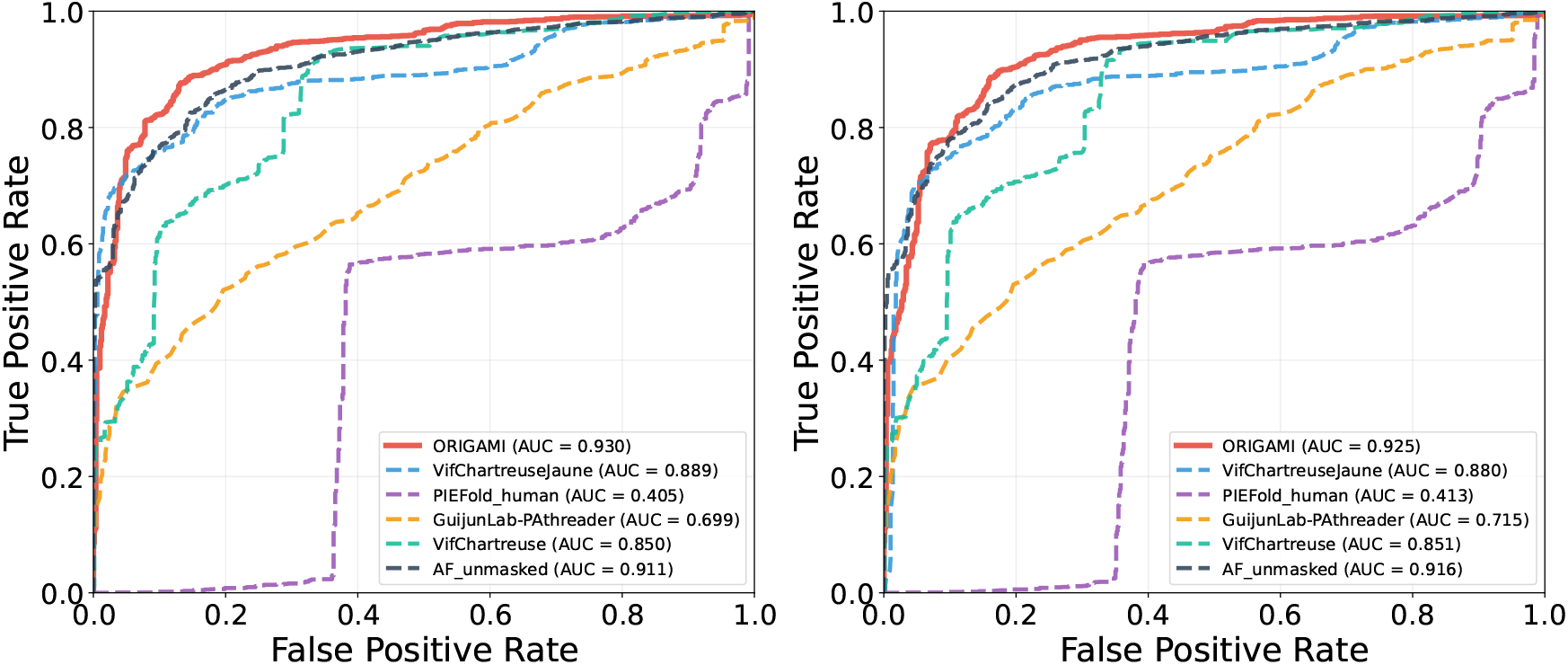
ROC curves for ORIGAMI and top CASP16 single-model EMA groups using iLDDT-based (left) and DockQ-based (right) model acceptability criteria.

**Figure 4** shows the correlations between ORIGAMI-predicted scores and the ground truth iLDDT and DockQ across the 12 CASP16 dimeric targets. Despite training exclusively on superposition-free iLDDT scores, ORIGAMI accurately reproduces superposition-based DockQ scores; and achieves Pearson correlations of 0.778 (iLDDT) and 0.729 (DockQ), with Spearman correlations of 0.760 and 0.693, and MAEs of 0.174 and 0.191, respectively.

**Figure 4.**
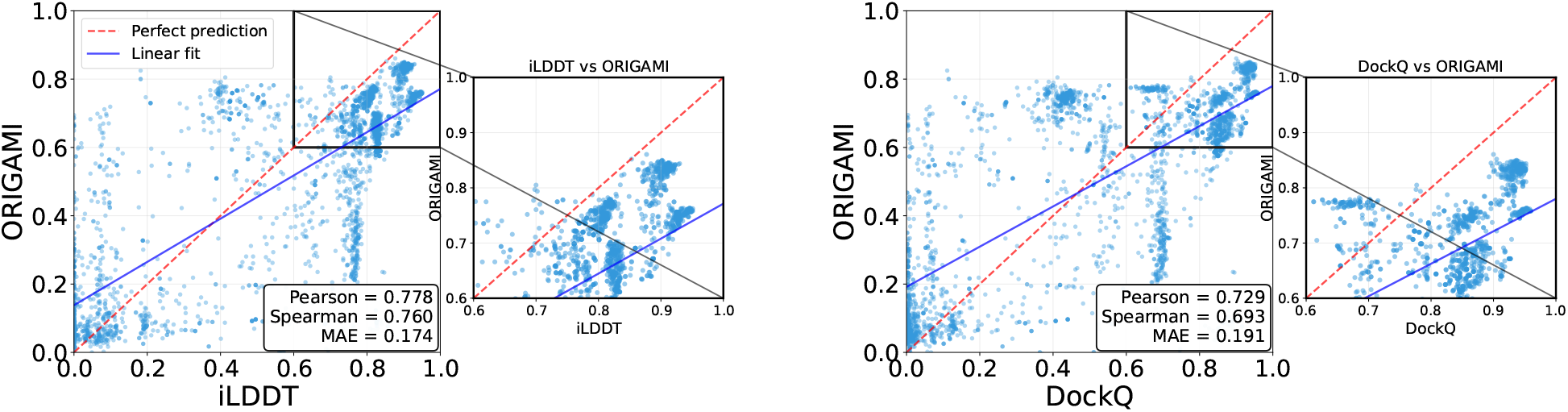
Comparison of ORIGAMI-predicted scores and ground truth iLDDT (left) and DockQ scores (right) on the CASP16 dimeric targets.

### 3.2 Performance on the CASP15 dataset

To further validate ORIGAMI’s generalizability, we evaluated its performance on a fixed benchmark comprising 24 protein complex targets from CASP15 (34). All methods, including ORIGAMI, three in-house methods, and nine top-performing CASP15 predictors, were evaluated on the same set of 5,264 dimeric models derived from these targets. **Figure 5** shows the hit rate analysis comparing ORIGAMI with three in-house deep learning methods and nine CASP15 (34) single-model EMA methods. The hit rate measures the fraction of target complexes for which a method successfully selects at least one high-quality model (iLDDT ≥ 0.236) within the top-*N* ranked predictions. At Top-1, APOLLO, DProQA (29), Guijunlab-RocketX, and VoroIF achieve the highest hit rate of 0.750, while ORIGAMI and VoroIFGNN (27) obtain 0.708. At Top-5, ORIGAMI achieves the highest hit rate of 0.917, followed by Guijunlab-RocketX at 0.875 and VoroIFGNN at 0.875, while the fourth best method, APOLLO, achieves 0.833. At Top-10, ORIGAMI, VoroIFGNN, and VoroIF all achieve 0.917. Among the four deep learning methods, DeepRank gnn esm (28) shows the lowest hit rates across all ranking thresholds, achieving 0.542, 0.583, and 0.833 at Top-1, Top-5, and Top-10, respectively. ORIGAMI demonstrates strong performance particularly at Top-5 and beyond, consistently ranking among the top-performing methods for protein complex model quality assessment on the CASP15 dataset. Detailed numerical results are provided in Supplementary Table 4.

**Figure 5.**
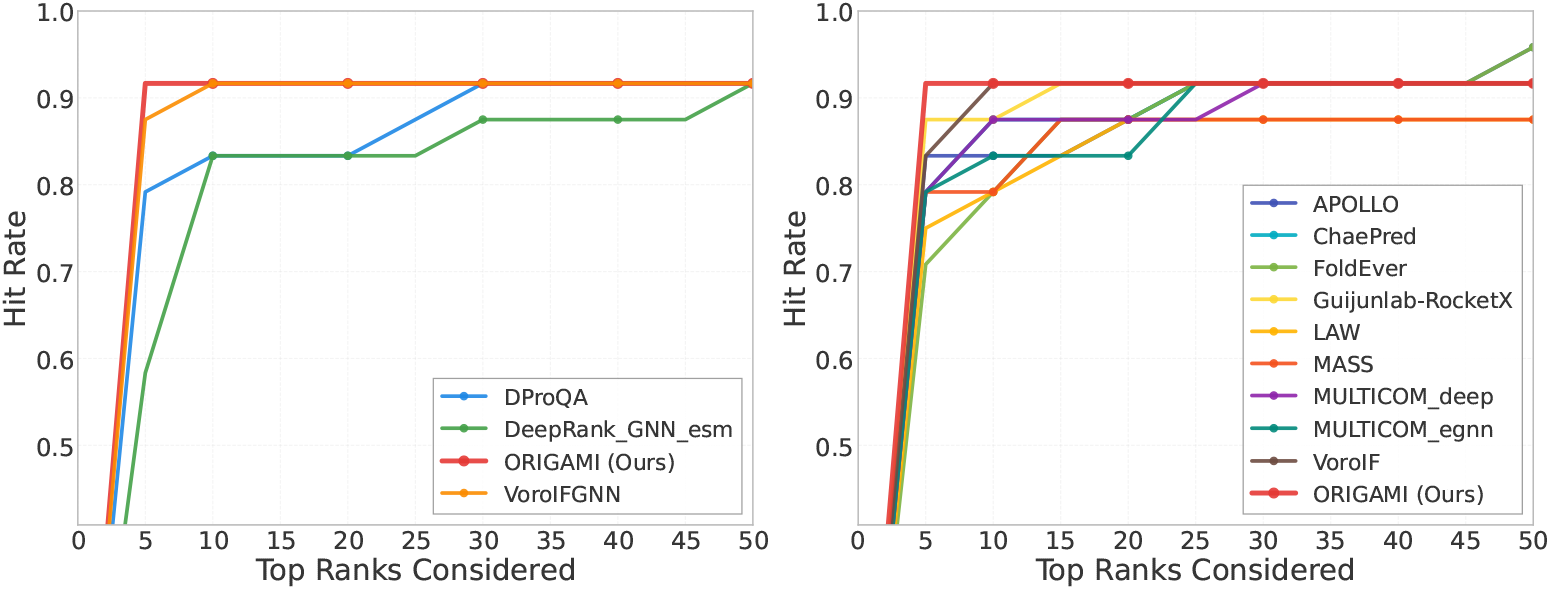
Performance on the CASP15 dataset. ORIGAMI was compared with three in-house deep learning methods (left) and nine top-performing CASP15 predictors (right). The figure shows the fraction of target complexes in the CASP15 dataset for which a method selected at least one acceptable model (within the top-*N* scored models).

### 3.3 Case study of protein–protein interface dynamics

Biological protein–protein interfaces often undergo distinct conformational substates during molecular motion. To evaluate whether ORIGAMI can capture such dynamic interface-level changes, we analyzed a 300 ns GROMACS molecular dynamics trajectory of the 1BRS barnase–barstar complex. It was previously reported that (53), for 1BRS, the fraction of preserved initial interface contacts remained around 90% during the first 125 ns and decreased to around 75% for the remainder of the simulation; and Jaccard-index-based clustering further separated the trajectory into two long-lived interface substates, corresponding approximately to 0–125 ns and 125–300 ns. We therefore used 125 ns as the reported interface substate transition point and compared ORIGAMI interface scores with an independent contact-preservation score over the trajectory. Weighted GDT-TS was included as a global structural reference that captures the global fold-level conformational stability of the individual chains weighted by their chain lengths.

As shown in Fig. 6A, the weighted GDT-TS score remains relatively stable throughout the trajectory, suggesting that the global fold-level conformational stability of the individual chains is largely preserved. In contrast, the contact-preservation score drops around the reported 125 ns transition, indicating an interface-level rearrangement rather than global structural degradation. Segment-averaged scores before and after the 125 ns transition (dotted lines in Fig. 6A) are contact preservation = 0.892 and 0.737, and ORIGAMI interface scores = 0.425 and 0.292, respectively. ORIGAMI follows this interface-level change, showing a Spearman correlation of *ρ* = 0.64 with contact preservation. The representative snapshots further illustrate this transition. At 115 ns, a local interface segment remains folded and compact relative to the native structure (Fig. 6B), with contact preservation = 0.827 and ORIGAMI score = 0.412. By 130 ns, the same region appears opened or partially unfolded at the interface (Fig. 6C), accompanied by a decrease in contact preservation to 0.692 and in ORIGAMI score to 0.298. These results suggest that ORIGAMI is quite sensitive to dynamic interface rearrangements that are not fully reflected by global fold-level similarity metrics.

**Figure 6.**
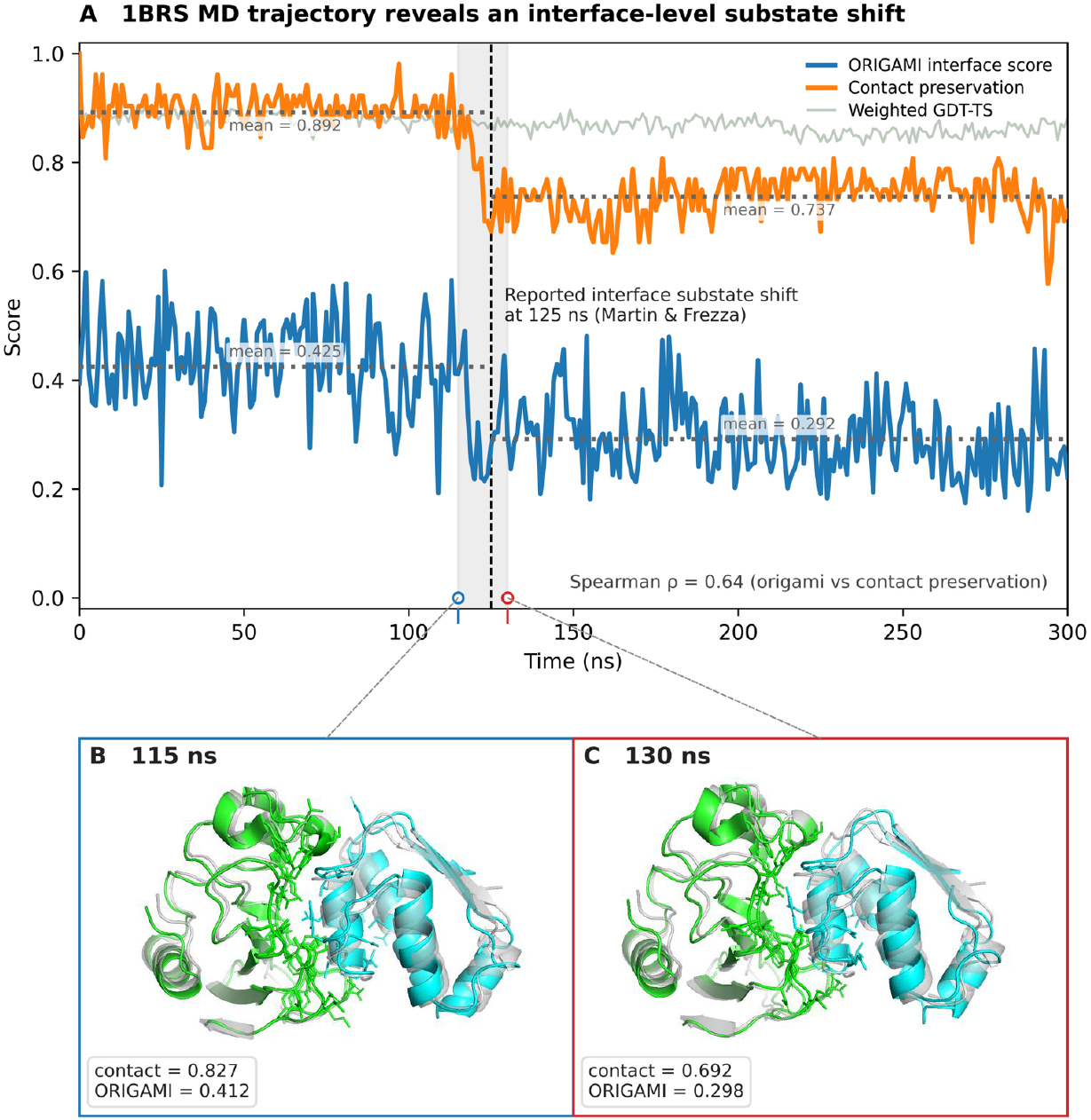
Dynamic interface assessment on the 1BRS MD trajectory. **(A)** ORIGAMI interface score, contact preservation, and weighted GDT-TS over the 300 ns MD trajectory. The shaded window marks 115–130 ns, surrounding the reported 125 ns interface substate shift. **(B, C)** Representative snapshots before and after the transition at 115 ns and 130 ns, respectively, overlaid with the native structure in semi-transparent gray. Interface residues are highlighted as sticks and colored according to their corresponding chains. The local interface region changes from a folded and compact state to an opened or partially unfolded state, accompanied by decreases in contact preservation from 0.827 to 0.692 and ORIGAMI score from 0.412 to 0.298.

### 3.4 Ablation study

To examine which components contribute to ORIGAMI’s performance, we performed a focused ablation study summarized in **Table 5**. The ablation study was organized into three parts: the overall architecture, the message-passing scheme, and the nearest-neighbor graph construction. Performance was assessed using Pearson correlation (*r*), Spearman correlation (*ρ*), and mean absolute error (MAE).

**Table 5.** Ablation study on the VoroIFGNN test dataset.

| Parameter | Value | Correlation |  | Error |
| --- | --- | --- | --- | --- |
| | | $r \uparrow$ | $\rho \uparrow$ | MAE ↓ |
| Architecture | **ORIGAMI** | **0.640** | **0.620** | **0.173** |
|  | SE(3)-Transformer | 0.488 | 0.457 | 0.203 |
|  | EGNN | 0.536 | 0.534 | 0.190 |
|  | Equiformer | 0.337 | 0.341 | 0.234 |
| Message-passing scheme | **GNN_SVP** | **0.640** | 0.620 | **0.173** |
|  | GNN_GVP | 0.626 | 0.603 | 0.173 |
|  | GIN_SVP | 0.613 | 0.602 | 0.191 |
|  | GIN_GVP | 0.638 | **0.624** | 0.176 |
|  | GAT_GVP | 0.592 | 0.581 | 0.184 |
| Nearest neighbor | Fixed-$k = 30$ | 0.627 | 0.602 | 0.174 |
|  | **Fixed-$k = 40$** | **0.638** | **0.618** | **0.172** |
| | Fixed-$k = 50$ | 0.624 | 0.614 | 0.176 |
| | Adaptive-$k$ | 0.577 | 0.593 | 0.200 |

At the architecture level, we compared ORIGAMI with three established equivariant graph neural network baselines: SE(3)-Transformer, EGNN, and Equiformer. Architectural details are provided in Supplementary Material 1.5. All models were trained on the same training set, selected using the same validation set, and evaluated on the same VoroIFGNN test dataset using iLDDT as the ground-truth score. ORIGAMI achieves the best performance across all reported metrics, with *r* = 0.640, *ρ* = 0.620, and MAE = 0.173. Among the baseline models, EGNN performs best, but still obtains lower correlation and higher error (*r* = 0.536, *ρ* = 0.534, MAE = 0.190). SE(3)-Transformer achieves *r* = 0.488, *ρ* = 0.457, and MAE = 0.203, while Equiformer achieves *r* = 0.337, *ρ* = 0.341, and MAE = 0.234. These results indicate that ORIGAMI’s improvement cannot be attributed solely to the general use of equivariance. Instead, the consistent gains over existing equivariant architectures support the effectiveness of ORIGAMI’s directed weight perceptron, vector-valued weights, and cross-product geometric filters for interface quality assessment.

We next examined the effect of the message-passing scheme and geometric perceptron design. We compared variants based on different message-passing backbones, including GNN with mean aggregation, GIN with learnable weighted-sum aggregation, and GAT with attention-based aggregation. These backbones were combined with geometric perceptron variants, including the Geometric Vector Perceptron (GVP), which uses linear vector combinations, and the Scalar–Vector Perceptron (SVP), which employs directed weights enabling dot- and cross-product interactions between vector features. Among the tested message-passing schemes, GNN SVP achieves the best overall performance, with the highest Pearson correlation (*r* = 0.640) and lowest MAE (0.173). GIN GVP achieves the highest Spearman correlation (*ρ* = 0.624), but shows slightly lower Pearson correlation and higher MAE than GNN SVP. Overall, these results suggest that the GNN SVP design provides the most favorable balance between correlation and prediction error, while the differences among the top-performing message-passing schemes remain relatively small.

We further examined the effect of nearest-neighbor graph construction, which controls the number of spatially closest residues involved in message passing. Among the fixed-*k* settings, performance peaks at fixed-*k* = 40 (*r* = 0.638, *ρ* = 0.618, MAE = 0.172), while both smaller fixed-*k* = 30 and larger fixed-*k* = 50 neighborhoods lead to reduced accuracy. This indicates that a balanced local neighborhood is important for stable message passing and effective aggregation of orientation-aware geometric features.

We also evaluated an adaptive-*k* nearest-neighbor strategy in which the number of neighbors varies with protein size: *k* = 24 for proteins with 0–100 residues, *k* = 32 for proteins with 100–200 residues, and *k* = 48 for proteins with more than 200 residues. This adaptive setting was designed to provide a smaller local receptive field for short proteins and a broader neighborhood for larger proteins. However, adaptive-*k* achieves lower performance (*r* = 0.577, *ρ* = 0.593, MAE = 0.200) than all fixed-*k* settings. This suggests that simply varying the neighborhood size according to protein length does not improve interface quality prediction on this benchmark and may introduce less relevant residues into the local interface representation. Overall, the fixed-*k* setting, especially fixed-*k* = 40, provides a more stable balance between capturing sufficient geometric context and preserving locality around the interface. Additional hyperparameters are provided in Supplementary Table 2.1.

## 4 Conclusion

We present ORIGAMI, an orientation-aware graph neural network for protein complex quality assessment that leverages SO(3)-equivariant geometric operations to capture fine-grained orientational relationships at protein-protein interfaces. By incorporating directional information beyond scalar-only representations, ORIGAMI effectively learns intricate geometric features that distinguish high-quality multimeric models from incorrect predictions. Our comprehensive evaluation demonstrates that ORIGAMI achieves competitive or superior performance across multiple interface quality assessment benchmarks, with particularly strong gains in the expanded CASP16 interface-level evaluation and in controlled comparisons against both non-equivariant and equivariant graph neural network baselines. Despite being trained to estimate the superposition-free iLDDT score, ORIGAMI also shows robust cross-metric generalization by reproducing superposition-based DockQ scores with high fidelity. In addition, evaluation on a 300 ns GROMACS molecular dynamics trajectory of the 1BRS barnase–barstar complex shows that ORIGAMI follows the reported interface substate transition and correlates with interface contact preservation over time, suggesting sensitivity to dynamic protein–protein interface rearrangements. Future extensions could incorporate additional geometric descriptors or adapt the architecture for related tasks, such as protein-protein docking scoring or interface design validation. Additional future directions include extending ORIGAMI to broader dynamic ensembles, heterogeneous chemical mediators such as water molecules, metal ions, and ligands, and large assemblies where long-range allosteric or many-body effects may influence interface quality.

## 5 Acknowledgements

This work was partially supported by the National Institute of General Medical Sciences (2R35GM138146 to D.B.) and the National Science Foundation (DBI2208679 to D.B.).

## 6 Data and Software Availability

The source code, trained models, and detailed instructions for using ORIGAMI are available at https://github.com/Bhattacharya-Lab/ORIGAMI. The training, validation and test datasets are available at https://zenodo.org/records/19076236.

## Notes

### Competing Interest Statement

The authors have declared no competing interest.

