## Supplementary Information for "ORIGAMI: Orientation-Aware Graph Neural Network for Assessing Multimeric Interfaces of Protein Complex Structures"

### 1 Methods

#### 1.1 Targets from CASP15 and CASP16

We collected multimer predictions from the CASP16 and CASP15 websites [1, 2]. For CASP16, we evaluated 39 multimeric targets, including both dimeric and higher-order oligomeric assemblies. To enable consistent interface-level assessment across complexes with different stoichiometries, each target was evaluated using reference-defined contacting subunit-pair interfaces. Specifically, interacting subunit pairs were identified from the experimental assembly, and all submitted models for that target were evaluated on the same set of contacting interfaces. This produced 60,314 model–interface instances from 12,916 models.

In addition to this full CASP16 multimer evaluation set, we considered a dimeric CASP16 subset for comparison with top CASP16 predictors. This subset consists of the common set of dimeric protein complexes predicted by all five top-performing predictors, comprising 12 targets and 3,000 models.

For CASP15, we collected the common set of dimeric protein complexes predicted by the three in-house methods and 10 top-performing CASP15 predictors, comprising 24 targets and 5,264 models. The full lists of targets used in this study are provided in **Table 1**.

**Table 1.** List of CASP targets used in the paper.

| **Data Set** | **Target List** |
|---|---|
| CASP15 ($n = 24$) | H1129, H1134, H1135, H1140, H1142, H1143, H1144, H1151, H1171, H1172, T1109, T1110, T1113, T1121, T1123, T1124, T1127, T1152, T1153, T1160, T1161, T1176, T1178, T1179 |
| CASP16_multimers ($n = 39$) | H1202, H1204, H1208, H1213, H1215, H1217, H1220, H1222, H1223, H1225, H1227, H1232, H1233, H1236, H1244, H1245, H1258, H1265, H1267, H1272, T1201o, T1206o, T1218o, T1219v1o, T1234o, T1235o, T1237o, T1240o, T1249v1o, T1249v2o, T1257o, T1259o, T1269v1o, T1270o, T1292o, T1294v1o, T1294v2o, T1295o, T1298o |
| CASP16_dimers ($n = 12$) | H1215, H1227, H1245, T1201, T1206, T1218, T1219v1, T1269v1, T1292, T1294v1, T1294v2, T1298 |

#### 1.2 Training, validation and test Datasets

**Voro-CASP datasets.** We constructed comprehensive datasets by combining protein complex models generated by VoroIFGNN [3] with structures from CASP13 [4] and CASP14 [5] competitions. The VoroIFGNN dataset employs a systematic model selection strategy based on VoroMQA-energy [6] and interface CAD-score [7] metrics. Specifically, the selection process operates in two stages: First, for each target's ranked list ordered by VoroMQA-energy score, models are selected such that their interface CAD-score values differ by 0.1 or more. Second, the ranked list is restricted to models with interface CAD-score equal to 0, and models are selected such that their binding site CAD-score values differ by 0.1 or more, again starting from the best VoroMQA-energy score. This systematic approach yielded up to 20 models per target (12.8 models on average), covering a wide range of accuracies in both detailed interface accuracy and binding-site accuracy. The selected diverse models, while predominantly incorrect, maintain better-than-random VoroMQA-energy values, creating deceptive decoys suitable for rigorous accuracy estimation method evaluation.

**Voro-CASP training datasets.** The training partition comprises 1,090 targets with 15,279 models, generated using the systematic selection protocol described above. This extensive training set provides comprehensive coverage of interface geometries and accuracy levels necessary for robust model learning, with identical generation protocols ensuring consistency across the dataset.

**Voro-CASP validation datasets.** The validation set contains 224 targets with 2,671 models, following the same rigorous selection methodology as the training data. This partition enables hyperparameter tuning and model selection while maintaining the same diversity in interface accuracy and binding-site accuracy as other dataset partitions.

**Voro-CASP testing datasets.** The testing partition includes 227 targets with 2,753 models, providing an independent evaluation benchmark. The testing set maintains the same generation protocols and model selection criteria, ensuring no target overlap between different evaluation phases while preserving the challenging nature of deceptive decoys for comprehensive method assessment.

**CASP15 and CASP16 datasets.** We collected multimer predictions from the CASP16 and CASP15 websites [1, 2]. For CASP16, we evaluated 39 multimeric targets, including both dimeric and higher-order oligomeric assemblies, using reference-defined contacting subunit-pair interfaces. Interacting subunit pairs were identified from the experimental assemblies, and all submitted models were evaluated on the same set of contacting interfaces for each target, yielding 60,314 model–interface instances from 12,916 models. We additionally considered a CASP16 dimeric subset consisting of the common set of dimeric protein complexes predicted by the five top-performing predictors, comprising 12 targets and 3,000 models. For CASP15, we evaluated 24 dimeric targets and 5,264 models from 10 top-performing predictors and the three baseline methods.

#### 1.3 Interface Local Distance Difference Test (iLDDT)

The accuracy of protein-protein interfaces was evaluated using the interface Local Distance Difference Test (iLDDT), a superposition-free metric that quantifies how well local distance relationships are preserved between atoms across different chains in the interface [8, 9]. The iLDDT score extends the standard LDDT methodology to specifically assess inter-chain atomic interactions, making it particularly suitable for evaluating the quality of protein complex predictions and binding interfaces.

For each atom pair $(i, j)$ where atoms $i$ and $j$ belong to different chains and are within an inclusion radius $R_0 = 15$ Å, the distance difference is calculated as:

$$\Delta d_{ij} = |d_{ij}^{\text{model}} - d_{ij}^{\text{reference}}| \tag{1}$$

where $d_{ij}^{\text{model}}$ and $d_{ij}^{\text{reference}}$ are the inter-atomic distances in the predicted model and reference structure, respectively. The preservation of distance relationships is evaluated using four tolerance thresholds: $t_1 = 0.5$ Å, $t_2 = 1.0$ Å, $t_3 = 2.0$ Å, and $t_4 = 4.0$ Å. For each threshold $t_k$, a distance is considered preserved if $\Delta d_{ij} \leq t_k$.

The iLDDT score is computed as:

$$\text{iLDDT} = \frac{1}{4} \sum_{k=1}^{4} \frac{1}{N} \sum_{(i,j) \in L} \mathbf{1}(\Delta d_{ij} \leq t_k) \tag{2}$$

where $N$ is the total number of inter-chain atom pairs within the inclusion radius, $L$ represents the set of all such atom pairs, and $\mathbf{1}(\cdot)$ is the indicator function that equals 1 if the condition is satisfied and 0 otherwise. The final iLDDT score ranges from 0 to 1, where higher values indicate better preservation of interface geometry and stronger agreement between the predicted and reference structures. In this study, protein LDDT scores (both intra-chain and interface) were calculated with an inclusion radius of 15 Å, accounting for the larger scale of protein-protein interactions and ensuring comprehensive assessment of interface accuracy.

#### 1.4 ORIGAMI architecture details

The ORIGAMI framework implements a multi-layer graph convolutional architecture that processes both scalar and vector node representations derived from interfacial residue features. The pipeline begins by analyzing 3D protein complex structures to locate and isolate interfacial regions where protein chains interact. Interfacial zones are defined as regions containing residues with at least one atom within 24 Å of an atom from a different chain. The extracted interfacial region is then modeled as a spatial $k$-nearest neighbor (k-NN) graph $G = (V, E)$, where each interfacial residue corresponds to a node $u \in V$ and each node is connected to its $k = 40$ spatially closest neighboring residues in $\mathbb{R}^3$.

Node representations use scalar–vector tuples $h_u = (s_u, V_u)$ with $s_u \in \mathbb{R}^C$ and $V_u \in \mathbb{R}^{C\times 3}$, while edges $E = \{e_{ij}\}_{i\neq j}$ between residues $i$ and $j$ from different protein chains are encoded with multi-dimensional scalar–vector tuples. The framework supports configurable graph convolution backbones, including standard GNN layers, Graph Attention Networks (GAT), and Graph Isomorphism Networks (GIN), enabling flexible adaptation to different protein complex assessment settings.

The resulting scalar and vector features are processed by six consecutively stacked SVGraphConv layers, each of which incorporates the core architectural unit, the Directed Weight Perceptron (DWP). Every DWP module consists of three sequential stages : (a) a *Directed Linear* stage that applies four geometric transformation pathways to scalar and vector inputs using directed weight parameters; (b) a *Non-Linearity* stage that applies ReLU activations to scalar components and sigmoid-based gating to vector components; and (c) a *Directed Interaction* stage that enables bidirectional information flow between scalar and vector feature modalities.

Each graph convolution layer implements an equivariant message-passing mechanism that aggregates information from k-nearest neighbors while preserving geometric relationships. The layer architecture includes message aggregation over spatial neighborhoods, layer normalization for training stability, scalar–vector perceptron feed-forward networks with activation functions tailored to dual-modality features, and dropout regularization to reduce overfitting during complex structure learning. ORIGAMI preserves SO(3)-equivariance through an equivariant message propagation scheme that performs coordinate transformations between local reference frames defined by residue backbone geometries throughout neighborhood aggregation.

After the final convolution layer, node-level representations are aggregated by scatter mean pooling to obtain a fixed-dimensional graph embedding capturing global interface characteristics. This graph-level representation is then passed through a readout module composed of sequential linear transformations with ReLU activation and dropout regularization, ultimately producing a single scalar output corresponding to the interface Local Distance Difference Test (iLDDT) score [9]. The predicted iLDDT value lies in $[0, 1]$ and provides a continuous estimate of interface binding quality and structural stability.

#### 1.5 Alternative architecture details

We evaluated several alternative equivariant architectures as additional baselines, including SE(3)-Transformer, Equiformer, and EGNN. All models were trained on the same train/validation split, consisting of 15,279 training examples and 2,671 validation examples. Unless otherwise specified, models were trained for 100 epochs using the AdamW optimizer with a weight decay of $10^{-6}$ and an L1/MAE loss objective. Accordingly, the reported training MAE, validation MAE, and validation loss all correspond to mean absolute error.

**SE(3)-Transformer.** The SE(3)-Transformer baseline was trained for 100 epochs with a learning rate of $10^{-4}$ using the AdamW optimizer and a weight decay of $10^{-6}$. The model was optimized using L1 loss, and the reported training MAE, validation MAE, and validation loss correspond to MAE/L1. Training used a batch size of 2, 2 data-loading workers, and a single GPU specified by `CUDA_VISIBLE_DEVICES=1`. Structures were truncated or filtered using a maximum residue limit of 1,200 residues, and graphs were constructed with a distance cutoff of 10.0 Å. The SE(3)-Transformer architecture used 2 layers, 2 attention heads, 3 degrees, and 8 channels.

**Equiformer.** The Equiformer baseline was implemented as a Graph Attention Transformer and trained for 100 epochs with a learning rate of $10^{-4}$ using AdamW and a weight decay of $10^{-6}$. The loss function was L1 loss, equivalent to MAE. Training was distributed across 4 GPUs using `torch.distributed.run --nproc_per_node=4`, with a batch size of 1 per GPU and 2 data-loading workers. Graphs were constructed using a 10.0 Å distance cutoff. The model used 4 layers, 64 basis functions, and 4 attention heads.

**EGNN.** The EGNN baseline was trained for 100 epochs with a learning rate of $10^{-5}$ using AdamW and a weight decay of $10^{-6}$. The model was optimized with L1 loss/MAE. Training used a batch size of 2, 2 data-loading workers, and a single GPU specified by `CUDA_VISIBLE_DEVICES=1`. Structures were truncated or filtered using a maximum residue limit of 800 residues, and graphs were constructed with a 10.0 Å distance cutoff. The EGNN model was implemented as an `EGNNRegressor` with a hidden dimension of 128 and 4 layers. Gradient clipping with a maximum norm of 1.0 was applied to improve numerical stability and prevent NaN values during training.

##### 1.6 Evaluation metrics

To assess ranking quality, we compute **Pearson and Spearman correlation coefficients** between predicted and actual lDDT scores. The **Pearson correlation coefficient** measures linear relationships and is defined as:

$$r = \frac{\sum_{i=1}^{n}(x_i - \bar{x})(y_i - \bar{y})}{\sqrt{\sum_{i=1}^{n}(x_i - \bar{x})^2}\sqrt{\sum_{i=1}^{n}(y_i - \bar{y})^2}}, \tag{3}$$

where $x_i$ and $y_i$ are the predicted and true lDDT scores, respectively, and $\bar{x}$ and $\bar{y}$ are their corresponding means.

The **Spearman rank correlation coefficient** evaluates monotonic relationships and is defined as:

$$\rho = 1 - \frac{6\sum_{i=1}^{n} d_i^2}{n(n^2 - 1)}, \tag{4}$$

where $d_i$ is the difference between the ranks of corresponding predicted and true lDDT scores.

Additionally, we calculate the **Mean Absolute Error (MAE)** as:

$$\text{MAE} = \frac{1}{n}\sum_{i=1}^{n} |y_i - \hat{y}_i|, \tag{5}$$

where $y_i$ represents the true lDDT score and $\hat{y}_i$ represents the predicted score for the $i$-th sample.

To evaluate the ability of each method to select high-quality models, we also compute a per-target **ranking loss**. For a given target $t$, let $D_t$ denote the set of decoys, $q_{tj}$ denote the true quality score of decoy $j$ measured by DockQ or iLDDT, and $\hat{q}_{tj}$ denote the score predicted by the ranking method. The best available decoy for target $t$ is defined as

$$j_t^* = \arg\max_{j \in D_t} q_{tj}, \tag{6}$$

whereas the top-ranked decoy selected by the method is

$$\hat{j}_t = \arg\max_{j \in D_t} \hat{q}_{tj}. \tag{7}$$

The per-target ranking loss is then defined as the difference between the DockQ or iLDDT score of the target's best decoy and the DockQ or iLDDT score of the top decoy selected by the ranking method:

$$\mathcal{L}_{\text{rank}}(t) = q_{tj_t^*} - q_{t\hat{j}_t}. \tag{8}$$

A ranking loss of zero indicates that the method successfully selected the best available decoy for that target, whereas larger values indicate a greater loss in model quality due to suboptimal ranking. The overall ranking loss across a benchmark containing $T$ targets is computed as:

$$\mathcal{L}_{\text{rank}} = \frac{1}{T}\sum_{t=1}^{T} \mathcal{L}_{\text{rank}}(t). \tag{9}$$

### 2 Results

#### 2.1 Comprehensive Ablation Study on VoroIFGNN_test Dataset

**Table 2.** Comprehensive Ablation Study on VoroIFGNN_test Dataset

| Parameter | Value | Correlation ↑ | | | Error ↓ | | Score ↑ |
|---|---|---|---|---|---|---|---|
| | | **Pearson** | **Kendall** | **Spearman** | **Ranking Loss** | **MAE** | |
| **Distance** | 20 | 0.608 | 0.423 | 0.596 | 0.3412 | 0.1817 | 0.6731 |
| | 22 | 0.603 | 0.419 | 0.588 | 0.3371 | 0.1818 | 0.6726 |
| | **24** | **0.606** | **0.417** | **0.586** | **0.3098** | **0.1769** | **0.6832** |
| | 26 | 0.594 | 0.412 | 0.580 | 0.3192 | 0.1786 | 0.6770 |
| | 28 | 0.602 | 0.416 | 0.585 | 0.3248 | 0.1796 | 0.6766 |
| | 30 | 0.594 | 0.406 | 0.572 | 0.3341 | 0.1795 | 0.6701 |
| **Topk** | 30 | 0.627 | 0.428 | 0.602 | 0.3162 | 0.1744 | 0.6872 |
| | 35 | 0.632 | 0.433 | 0.611 | 0.3209 | 0.1704 | 0.6891 |
| | **40** | **0.638** | **0.440** | **0.618** | **0.3095** | **0.1719** | **0.6946** |
| | 45 | 0.631 | 0.434 | 0.609 | 0.3136 | 0.1742 | 0.6901 |
| | 50 | 0.624 | 0.437 | 0.614 | 0.3049 | 0.1763 | 0.6924 |
| **Num_layers** | 5 | 0.626 | 0.429 | 0.605 | 0.3260 | 0.1739 | 0.6845 |
| | **6** | **0.640** | **0.440** | **0.620** | **0.3131** | **0.1733** | **0.6934** |
| | 7 | 0.629 | 0.429 | 0.618 | 0.3132 | 0.1750 | 0.6902 |
| | 8 | 0.629 | 0.430 | 0.608 | 0.3135 | 0.1779 | 0.6881 |
| **Models** | **Gnn_svp** | **0.640** | **0.440** | **0.620** | **0.3131** | **0.1733** | **0.6934** |
| | Gnn_gvp | 0.626 | 0.432 | 0.603 | 0.3167 | 0.1734 | 0.6879 |
| | Gin_svp | 0.613 | 0.428 | 0.602 | 0.3275 | 0.1911 | 0.6764 |
| | Gin_gvp | 0.638 | 0.445 | 0.624 | 0.3023 | 0.1759 | 0.6969 |
| | Gat_gvp | 0.592 | 0.411 | 0.581 | 0.3090 | 0.1843 | 0.6782 |
| **Vector_hid** | (1,1) | 0.623 | 0.431 | 0.609 | 0.3193 | 0.1783 | 0.6856 |
| | (8,8) | 0.638 | 0.441 | 0.621 | 0.3178 | 0.1738 | 0.6917 |
| | **(32,16)** | **0.640** | **0.440** | **0.620** | **0.3131** | **0.1733** | **0.6934** |

#### 2.2 CASP15 Hit Rate Table

**Table 3.** Hit Rate Analysis for CASP15 (iLDDT $\geq$ 0.236) . All methods were evaluated on identical 24 targets and 5,264 models to ensure fair comparison.

| **Method** | **Type** | **Top-1** | **Top-5** | **Top-10** | **Top-20** | **Top-50** |
|---|---|---|---|---|---|---|
| DProQA | in-house | 0.750 | 0.792 | 0.833 | 0.833 | 0.917 |
| DeepRank_GNN_esm | in-house | 0.542 | 0.583 | 0.833 | 0.833 | 0.917 |
| **ORIGAMI** (Ours) | in-house | 0.708 | 0.917 | 0.917 | 0.917 | 0.917 |
| VoroIFGNN | in-house | 0.708 | 0.875 | 0.917 | 0.917 | 0.917 |
| APOLLO | CASP | 0.750 | 0.833 | 0.833 | 0.875 | 0.958 |
| ChaePred | CASP | 0.583 | 0.792 | 0.875 | 0.875 | 0.917 |
| FoldEver | CASP | 0.625 | 0.708 | 0.792 | 0.875 | 0.958 |
| Guijunlab-RocketX | CASP | 0.750 | 0.875 | 0.875 | 0.917 | 0.917 |
| LAW | CASP | 0.500 | 0.750 | 0.792 | 0.875 | 0.875 |
| MASS | CASP | 0.583 | 0.792 | 0.792 | 0.875 | 0.875 |
| MULTICOM_deep | CASP | 0.458 | 0.792 | 0.875 | 0.875 | 0.917 |
| MULTICOM_egnn | CASP | 0.708 | 0.792 | 0.833 | 0.833 | 0.917 |
| VoroIF | CASP | 0.750 | 0.833 | 0.917 | 0.917 | 0.917 |
